# MDTOMO: Continuous conformational variability analysis in cryo electron subtomogram data using flexible fitting based on Molecular Dynamics simulations

**DOI:** 10.1101/2023.02.25.529934

**Authors:** Rémi Vuillemot, Isabelle Rouiller, Slavica Jonić

## Abstract

Cryo electron tomography (cryo-ET) allows observing macromolecular complexes in their native environment. The common routine of subtomogram averaging (STA) allows obtaining the three-dimensional (3D) structure of abundant macromolecular complexes, and can be coupled with discrete classification to reveal conformational heterogeneity of the sample. However, the number of complexes extracted from cryo-ET data is usually small, which restricts the discrete-classification results to a small number of enough populated states and, thus, results in a largely incomplete conformational landscape. Alternative approaches are currently being investigated to explore the continuity of the conformational landscapes that *in situ* cryo-ET studies could provide. In this article, we present MDTOMO, a method for analyzing continuous conformational variability in cryo-ET subtomograms based on Molecular Dynamics (MD) simulations. MDTOMO allows obtaining an atomic-scale model of conformational variability and the corresponding free-energy landscape, from a given set of cryo-ET subtomograms. The article presents the performance of MDTOMO on a synthetic ABC exporter dataset and an *in situ* SARS-CoV-2 spike dataset. MDTOMO allows analyzing dynamic properties of molecular complexes to understand their biological functions, which could also be useful for structure-based drug discovery.

## 1. Introduction

Cryo electron tomography (cryo-ET) is a structural biology technique that allows determining the three-dimensional (3D) structure of abundant biological macromolecular complexes. This technique is particularly important as it allows studying the complexes and their interactions *in situ*, i.e., in their native cellular environment^1^.

In cryo-ET, a series of images are recorded by titling the sample over a range of angles. During the image collection over multiple tilts, the sample accumulates radiation damage. The sample damage is reduced by (1) lowering the electron dose per image (the electron dose allowed to be accumulated by the sample is spread over the whole tilt range) and (2) collecting images over a smaller tilt range (usually between −60° and +60° in steps of 1° or 2°). The low electron dose to which the sample is exposed during the image collection results in images with a low signal-to-noise ratio (SNR), which in turn results in a noisy 3D volume (the so-called tomogram) that is obtained by 3D reconstruction from the tilt-image series. The limited tilt range used for the data collection results in an anisotropic resolution of the tomogram (the so-called missing-wedge problem)^2^. A macromolecular complex extracted from a tomogram into an individual volume (the so-called subtomogram) is difficult to analyze due to the low SNR and the missing-wedge-induced deformations. Therefore, many instances of the same complex extracted from a tomogram into subtomograms are usually iteratively aligned and averaged (the procedure referred to as subtomogram averaging, STA), which increases the SNR and reduces the missing-wedge-induced deformations.

STA can be combined with discrete 3D classification to tackle the conformational heterogeneity of the complexes^3–6^. The goal of the classification is to make the groups (classes) of subtomograms that are as homogeneous as possible regarding the conformation of the complex. To obtain a subtomogram average of good quality from a class, the class should be highly populated. However, considering a relatively small number of available subtomograms, discrete classification methods are usually used with a small number of classes, which makes them ill-suited in the context of highly heterogeneous datasets composed of a continuum of conformational states. This heterogeneity issue, known as continuous conformational variability, is well-known in single particle (SPA) cryo electron microscopy (cryo-EM) and is currently an active field of research^7–16,17^. The number of methods for analyzing continuous conformational variability in SPA datasets has been growing over the last few years, but only a few such methods have been proposed for cryo-ET so far^18,19^. This is likely due to the challenges of analyzing cryo-ET data that were mentioned earlier, namely low SNR, missing-wedge-induced deformations, and small number of subtomograms. Also, volumetric data processing induces additional computational complexity.

Recently, a method named Tomoflow has been proposed for analyzing continuous conformational variability in cryo-ET subtomograms, which uses a computer vision technique of dense 3D optical flow to estimate the displacement of voxels in each subtomogram with respect to the voxels in a global subtomogram average^18^. This subtomogram average (reference volume) is calculated at each iteration from all subtomograms that were elastically and rigid-body aligned with respect to the subtomogram average obtained in the previous iteration^18^. The reference volume used for the TomoFlow initialization can either be a subtomogram average obtained by classical STA from the given set of subtomograms or it can be a cryo-EM map. The estimated displacements are projected onto a linear subspace through principal component analysis (PCA) to place the subtomograms on a low-dimensional conformational landscape.

HEMNMA-3D is another method for analyzing continuous conformational variability in cryo-ET subtomograms, which is able to use either a reference-volume voxel displacement (as TomoFlow) or a reference-structure atomic displacement to match the conformational states in subtomograms^19^. The development of HEMNMA-3D was inspired by the SPA methods that describe the continuous conformational variability by displacing atoms of an available molecular structure (reference structure) to match the conformational states in the data^7,12,13,16^. Such methods restrain the conformational exploration to the energy region that preserves the structural elements of the complex, which reduces the risk of overfitting low-SNR data. They displace atoms of a reference structure using different methods to model the deformation field, such as normal mode analysis^7,12^, MD simulations^13,15^, and Zernike polynomials^16^. HEMNMA^7^ and its deep-learning-based extension DeepHEMNMA^12^ perform a flexible alignment of the reference structure to single particle images along the direction in the energy landscape that is defined by a set of vectors of collective motions, called normal modes. The mentioned HEMNMA-3D method^19^ is an extension of HEMNMA^7^ to normal-mode-based analysis of subtomograms. Another SPA method follows a direction in the energy landscape that is defined by Zernike polynomials^16^. However, the methods based on normal modes and Zernike polynomials restrict the estimation of the dynamics to a predefined direction in the energy landscape (the direction defined by normal modes or Zernike polynomials), which may not be sufficient to fully describe the conformational heterogeneity in the data. Contrary to these methods, a recently proposed SPA method named MDSPACE uses molecular dynamics (MD) simulations, in combination with normal modes, which allows a better exploration of the energy landscape^13^. For atomic displacements using a combination of normal modes and MD simulations, MDSPACE employs the algorithm named Normal Mode Molecular Dynamics (NMMD)^20^, which was initially proposed for flexible fitting of atomic structures into cryo-EM maps in the context of structural modeling.

In this article, we present MDTOMO, a novel approach for analyzing continuous conformational variability in subtomograms, which is based on extracting atomic models from subtomograms by flexible fitting of a reference atomic structure using a combination of normal modes and MD simulation. MDTOMO analyzes each individual subtomogram by performing NMMD-based flexible fitting to extract the atomic model that matches the conformational state in the subtomogram. MDTOMO allows interpreting datasets in the form of atomic conformational landscapes and their further analysis in terms of free-energy landscapes. Moreover, MDTOMO allows obtaining subtomogram averages from localized regions in the energy landscape. While other MD-based flexible fitting methods are used for fitting cryo-EM maps or SPA images, this is the first method that uses MD-based flexible fitting in the context of analyzing subtomograms.

We tested MDTOMO on a synthetic dataset of an ABC exporter^21^ and *in situ* dataset of a SARS-CoV-2 spike protein (S)^22^. The synthetic dataset provides a controlled environment where the results of the method can be compared with the ground-truth data, which allows assessing the method performance and robustness to noise and missing-wedge-induced deformations. Although MDTOMO does not currently correct for the missing wedge, it was able to produce meaningful results even for high amounts of noise. MDTOMO applied to *in situ* data indicates its ability to decipher the conformational landscape as a continuum of intermediate states.

## 2. Methods

MDTOMO is a method for extracting information about continuous conformational variability of macromolecular complexes from subtomograms. More precisely, by flexibly fitting an initial atomic model (reference structure) to each subtomogram, MDTOMO can estimate an atomic-scale model of the conformational variability landscape from which it is possible to obtain the energy landscape. Moreover, MDTOMO can visualize principal conformational changes in terms of the atomic displacements or the subtomogram averages of localized regions in the energy landscape.

First, the subtomograms are pre-aligned using a rigid-body alignment with respect to the initial model. Then, the conformational state in each subtomogram is estimated by performing “biased” NMMD simulations. NMMD flexibly displaces the initial atomic coordinates to match the underlying conformational state in the subtomogram. It extracts the conformational state and refines the rigid-body pre-alignment by estimating the rigid-body rearrangement that occurred during the simulation. Finally, the obtained set of atomic models is projected onto a low-dimensional space (representing an essential conformational space), which allows visualizing conformational distribution and deciphering potential trajectories of conformational changes.

To speed up the NMMD fitting of multiple subtomograms and reduce the risk of overfitting noise in the data, MDTOMO can be used with a coarse-grained initial atomic model, as shown in this article.

### Normal Mode Molecular Dynamics

To reduce the computational cost, MDTOMO uses the recently proposed Normal Mode Molecular Dynamics (NMMD)^20^ to handle the MD simulation. NMMD is a method that incorporates Normal Mode Analysis (NMA) into the simulation to increase the speed of the global-dynamics atomic displacements allowing a better efficiency of the fitting. In NMMD, a set of low-frequency normal modes, representing the most collective motions (the motions that move the largest number of atoms synergistically), is incorporated in the MD integration scheme to encourage the MD simulation to move along the normal mode directions. The normal modes allow a fast displacement along the global dynamics and the MD simulation ensures the structural stability of the system and corrects for the distortions that can be induced by normal modes. The incorporation of normal modes typically reduces the length of the simulation required to fit large-amplitude conformational changes and therefore allows reducing the overall simulation length and the computational cost of the simulation. NMMD has been used to fit atomic models into cryo-EM maps and proved to be, on average, 40% faster than the flexible fitting solely based on MD simulation^20^. In MDTOMO, NMMD is used to fit an initial atomic model into each subtomogram from the given set of subtomograms.

As in standard MD-based fitting approaches (solely based on MD simulation), NMMD incorporates an additive biasing potential to the total energy that is minimized during the fitting. This additional potential biases the simulation by attracting the atoms into the regions of highest densities in the given EM map while the MD simulation maintains the integrity of the structure. As in the majority of the fitting approaches^20,23–25^, the biasing potential in MDTOMO is based on the correlation coefficient (CC) between the experimental map and a synthetic map calculated from the atomic model that is being fitted.

### Coarse-grained simulations

To further reduce the computational cost of the analysis, we used a coarse-grained model based on off-lattice Cα Gō model^26^, which drastically reduces the computational time compared to all-atom simulations. The Gō-like models are a class of models that share similar concepts as the model proposed by Gō (e.g., defining the long-range interactions based on the native contacts of the initial model, which tends to bias the simulation towards the reference). Cα Gō model represents the protein with Cα atoms of the amino acid chain and captures better the native dynamics than the original Gō model^27^. Here, the coarse-grained models were obtained by the SMOG2 software^28^.

### Low-dimensional conformational space and free-energy calculation

As MDTOMO results in one atomic model per subtomogram, a further analysis must be performed to visualize the conformational landscape. In this context, the set of atomic models obtained by MDTOMO can be projected onto a low-dimensional (e.g., 2D or 3D) space. In this study, two methods were used to perform the dimension reduction of the atomic coordinates, namely Principal Component Analysis (PCA) and Uniform Manifold Approximation and Projection (UMAP)^29^. PCA is the well-established technique for dimension reduction, offering a fast, deterministic and linear reduction method, which is useful to describe the overall variability in most datasets. UMAP is a recent method that allows capturing the non-linear motions, which can, in some cases, achieve a better separation of the different populations compared to PCA. The density of points obtained in the low-dimensional space can then be converted into free energy differences through the Boltzmann factor Δ*G*/*k_B_ T* = –ln(*n*/*n*_0_) by counting the number of particles *n* in each region of the space and the number of particles in the most populated region *n*_0_, *k_B_* being the Boltzmann constant and *T* the temperature of the system. Here, *T* was set to 300 K.

### Rigid-body pre-alignment of subtomograms

MDTOMO refines the rigid-body pre-alignment of the subtomograms with the initial model. The pre-alignment prevents the simulation to get trapped into local minima. For the synthetic ABC exporter experiments, the pre-alignment was obtained by STA based on the fast rotational matching algorithm (FRM)^30^ implemented in Scipion^31^. Concerning the *in situ* dataset of SARS-CoV-2 spikes, we used the STA alignment that had been obtained in the original work^22^.

The rigid-body pre-alignment is refined in MDTOMO by estimating the rigid-body rearrangements that occurred during the MD simulation, and using them to correct the initial alignment. The estimation of the rigid-body alignment parameters is obtained by optimizing the rotation and the translations that minimize the root mean square deviation (RMSD) between the initial conformation and the fitted conformation, using an optimization algorithm based on singular value decomposition that is available in BioPython^32^.

### Software implementation

MDTOMO is implemented as part of ContinuousFlex^33^, a software package for continuous conformational heterogeneity analysis in SPA and cryo-ET data, which is also available as a plugin of Scipion^31^. The Scipion back-end allows a user-friendly graphical interface for MDTOMO and integrating the data analysis results from most of the recent cryo-ET methods through the new ScipionTomo suite^34^. The MD and NMMD simulation back-ends are handled by GENESIS^35^.

## 3. Results

In this section, we show the performance of MDTOMO using a set of synthetic subtomograms of an ABC exporter^21^ and using a set of experimental *in situ* subtomograms of a SARS-CoV-2 spike protein (S)^22^.

### Experiment with a synthetic dataset of ABC exporter

#### Synthetic dataset of ABC exporter

To test the performance of MDTOMO, we synthesized a dataset as close as possible to experimental cryo-ET conditions. The dataset simulates a highly heterogeneous cryo-ET dataset of a heterodimeric ABC exporter, TmrAB, along the substrate translocation cycle. During the translocation cycle, the ABC exporter changes its conformation from outward-facing to inward-facing, allowing the substrate to enter the intracellular cavity. ATP-binding first induces closing of the intracellular gate (an occluded conformation of the exporter, where both intracellular and extracellular gates are closed) and, then, opening of the extracellular gate (an outward-facing conformation, where the extracellular gate opens while the intracellular gate remains closed), which allows a release of the substrate in the extracellular environment. The structure of TmrAB in inward-facing, outward-facing, and occluded conformations were derived from cryo-EM maps^21^ and are available in the Protein Data Bank (PDB) under the accession codes 6RAF, 6RAH, and 6RAK, respectively. Figure 1a shows the PDB: 6RAK structure of TmrAB in the occluded conformation while Figure 1b shows a sketch of the translocation cycle, with all three mentioned conformational states of TmrAB during the cycle.

**Figure 1:**
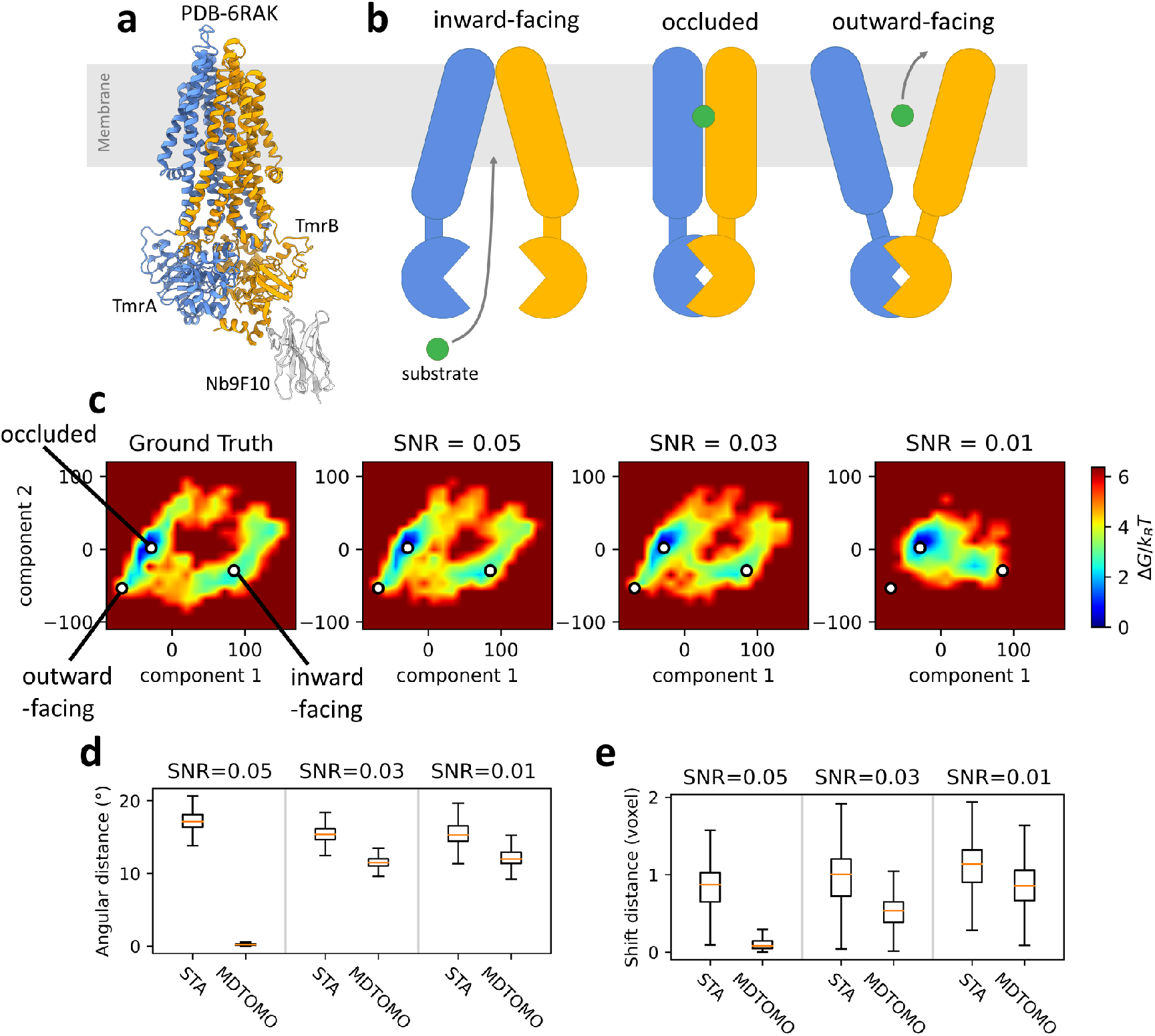
MDTOMO analysis of a synthetic dataset of TmrAB. (a) Structure of TmrAB in the occluded conformation. (b) Sketch of the substrate translocation process, from left to right: inward-facing, occluded, and outward-facing conformations of TmrAB. (c) Free-energy landscapes along the first two PCA components, from left to right: the ground-truth data used to synthesize the subtomograms and the MDTOMO results for the datasets with the SNR of 0.05, 0.03, and 0.01. The location of the three given PDB structures (6RAF, 6RAH, and 6RAK), used to target the MD simulations during the data synthesis, are also shown (white discs) in the free-energy landscapes. (d-e) Comparison of the initial and final rotational (d) and translational (e) alignment errors, for the three datasets (three SNR values).

The conformational heterogeneity was generated by standard MD-based flexible fitting (without normal mode acceleration) to obtain a dataset with realistic conformational variability, composed of a number of more populated conformational states (representing more stable states), close to the three mentioned states of TmrAB during the translocation cycle, and a number of less populated, intermediate states between these more stable states. The MD simulation, starting from the occluded conformation, was directed towards one of the two other conformational states at a time (inward-facing or outward-facing), alternating between these two states, by choosing the biasing potential that forces the simulation to go to one of the two conformational states. The biasing potential was defined by a simulated EM map of the corresponding conformational state which was changed every 10 picoseconds, resulting in a simulated MD trajectory of the exporter alternating between the outward-facing and inward-facing conformations while passing through a large number of intermediate states, including the occluded conformation. The MD simulation was performed for 1 nanosecond using Cα-Gō model. One can notice that a 10-picosecond period may not be long enough for reaching the target conformation for the fitting, meaning that the fitted model at the end of each 10-picosecond MD-based flexible fitting run may be slightly different from the target conformation. Nevertheless, the dataset consisting of a continuum of conformational states visited during this MD simulation is rich and representative of the translocation cycle enough for performing the tests of the method that is proposed here.

To simulate the sub-tomograms, we followed a previously published method^18,19^. We started by collecting 3000 atomic models from the simulated trajectory and we converted each model to a volume of size 100^3^ voxels and a voxel size of (2 Å)^3^ using a method based on scattering factors^36^. The obtained volumes were low-pass filtered at a cutoff frequency of 1/(6 Å) to take into account the impact of various imperfections (e.g., average radiation damage per image and tilt-series alignment errors). Then, the volumes were rotated and shifted randomly, and projected with a tilt angle from − 60° to +60° using an angular step of 2°. The tilt-series images were modulated by a contrast transfer function (CTF) that corresponds to a 200 kV microscope, a defocus of −0.5 μm, and a spherical aberration of 2 mm. Following a method that adds Gaussian noise before and after the CTF^37^, three sets of images were obtained corresponding to a SNR of 0.05, 0.03, and 0.01. The synthetic subtomograms were obtained from the simulated images using weighted back-projection in Fourier space^38^. The subtomograms were aligned using STA based on the FRM algorithm^30^ in Scipion.

#### Recovery of the ground-truth conformational space

We applied MDTOMO to each of the synthetic datasets (one for each SNR value) composed of 3000 subtomograms using 100-ps MD simulations starting from the PDB:6RAK model (occluded conformation). Figure 1c shows the free-energy landscape along the first two PCA components for the synthetic MD trajectory (ground truth) and the free-energy landscapes obtained with MDTOMO for each of the three datasets (subtomograms obtained from the images with the SNR of 0.05, 0.03, and 0.01). We observe that MDTOMO was able to recover the conformational landscape very accurately for the datasets with the SNR of 0.05 and 0.03. Indeed, Figure 1c shows that the most dense regions of the conformational space (i.e., the regions of the lowest free energy) are those that are the closest to the three given PDB structures (6RAF, 6RAH, and 6RAK) that were used to target the MD simulations during the data synthesis. In the case of the dataset with the highest level of noise (SNR of 0.01), the outward-facing conformation was detected less accurately (some densities are missing for the outward-facing state) than the two other states (occluded and inward-facing).

#### Recovery of the rigid-body alignments

The final rigid-body alignment, obtained by MDTOMO, was compared with the ground truth rigid-body displacement (the rotations and shifts used to synthesize the data). The initial rigid-body alignment, done by STA, was also compared with the ground truth rigid-body displacement. The initial and final alignment errors are compared in Figure 1d and Figure 1e, which show respectively the rotational and translational errors for the three datasets (three SNR values). We observe that, for both rotational and translational parameters and each value of the SNR, MDTOMO allows a refinement of the rigid-body alignment (the final errors are smaller than the initial errors).

### Experiment with a dataset of SARS-CoV-2 spike protein *in situ*

#### In situ dataset of SARS-CoV-2 spike protein

The SARS-CoV-2 spike (S) protein is a trimer of three identical glycoproteins (Figure 2a). It is involved in the cell infection as it mediates the viral entry by binding to the angiotensin-converting enzyme 2 (ACE2) receptor on the cell surface^39,40^. The S-protein head includes three receptor binding domains (RBDs) that are located at the top of the spike and exists in different conformations on the viral capsid. We used MDTOMO to analyze a set of experimental subtomograms of the S protein *in situ* that had previously been obtained and analyzed using STA and 3D classification (the corresponding tilt-series data are available in the EMPIAR database; accession code: EMPIAR-10453)^22^. In that previous publication^22^, the analysis of 20,830 subtomograms resulted in several classes, out of which two classes were further described. They correspond to two distinct conformational states, out of which one has three closed RBDs (fully closed state) and the other has one RBD opened (the classes containing 8,273 and 4,321 subtomograms, respectively).

**Figure 2:**
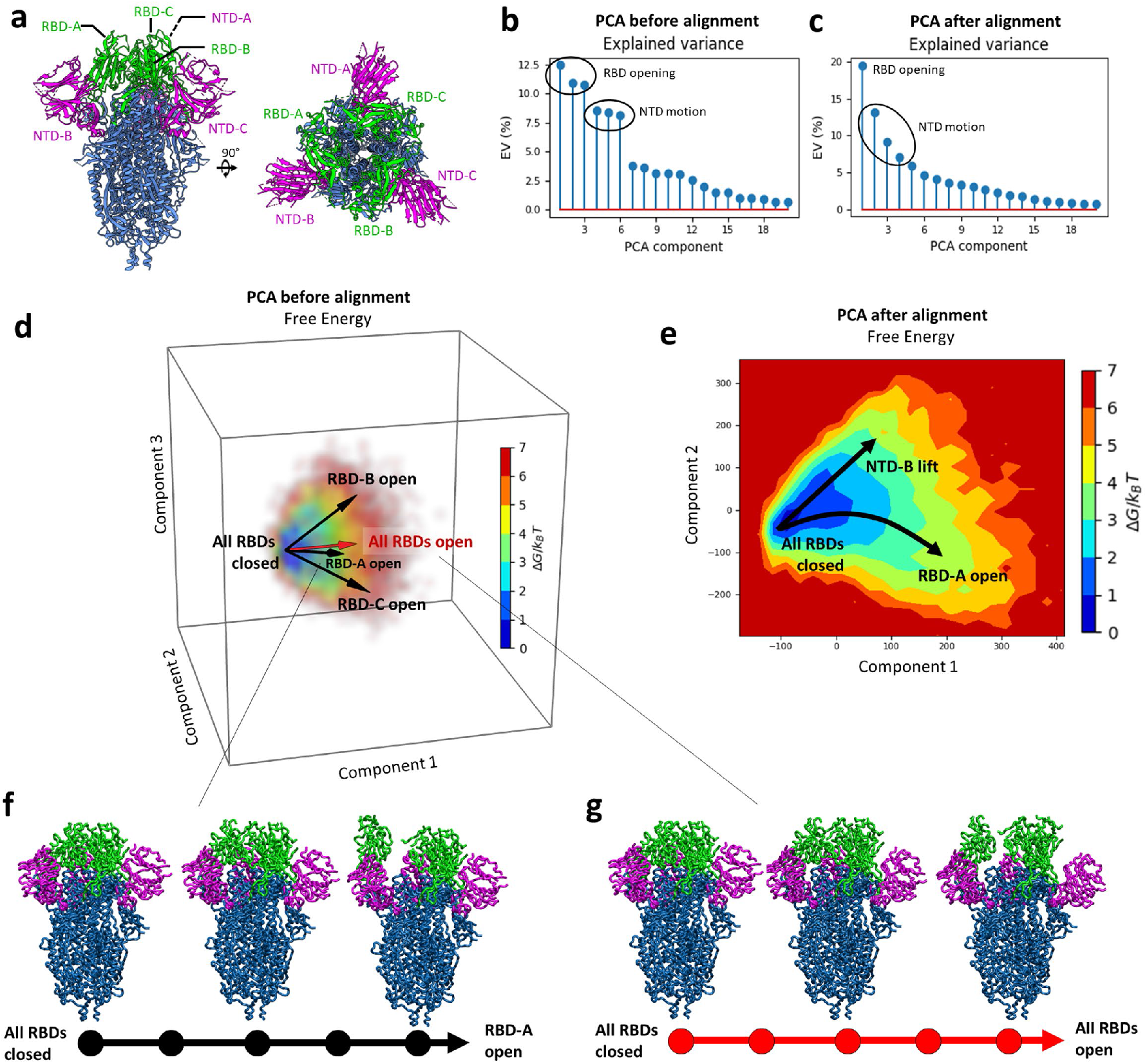
PCA analysis of MDTOMO results with a cryo-ET dataset of SARS-CoV-2 S protein. (a) Structure of the S protein (PDB 6VXX). (b-c) Explained variance of the PCA space obtained before (b) and after (c) reducing the number of principal components to describe opening of RBDs. (d) Free-energy landscape determined by the first three principal components, obtained before reducing the number of components to describe opening of RBDs. (e) Free-energy landscape determined by the first and second principal components obtained after reducing the number of components to describe opening of RBDs. The arrows show the directions associated with the RBD and NTD motions discussed here. (f) RBD-A opening trajectory following the direction “RBD-A open” shown in (d). (g) Trajectory of opening of all three RBDs together following the direction “All RBDs open” shown in (d).

MDTOMO was used to analyze the entire set of 20,830 subtomograms, which were first downsampled from their original size (256^3^ voxels) to the size of 128^3^ voxels and then pre-aligned using the rigid-body alignment parameters obtained by STA in the previous publication^22^. The initial conformation for the NMMD fitting with MDTOMO was PDB 6VXX, which corresponds to the fully closed state. The simulations included an initial energy minimization, followed by 100-ps NMMD simulations incorporating the 10 lowest frequency normal modes, with a force constant of 7000 kcal/mol.

#### Continuous conformational landscape of the S protein

The conformational variability was extracted by PCA from the set of atomic models obtained by MDTOMO. This PCA revealed multiple asymmetric conformations of the S protein. The plot in Figure 2b shows the variance described by each PCA component, and indicates that the first six principal components are responsible for more than 60% of the total variability. By exploring the conformational changes in different directions in the PCA space for the first six components, we could observe that the first three principal components (the most dominant motions) describe asymmetric opening and closing motions of the RBDs, whereas the next three principal components (less dominant motions) describe asymmetric up and down motions of the N-terminal domains (NTDs). Here, “asymmetric motion” means that each domain (RBD or NTD) can move independently from other domains.

Figure 2d shows a space determined by the first three principal components. Each of the three black arrows in this space, denoted by “RBD-A open”, “RBD-B open”, and “RBD-C open” (Figure 2d), shows a direction of motion that mainly describes a single RBD opening (one RBD is opening while the other two RBDs remain closed), for each of the three RBDs denoted in Figure 2a by “RBD-A”, “RBD-B” and “RBD-C”, respectively. Figure 2f shows the conformational change along the “RBD-A open” direction (opening of the RBD-A domain). In other directions of this space (Figure 2d), we could observe other conformational changes. In particular, when going from low-energy regions to high-energy regions in other directions than those indicated by black arrows (Figure 2d), we could observe opening of more than one RBD. For instance, the red arrow denoted by “All RBDs open” in Figure 2d indicates a direction along which all three RBDs are opening together, and Figure 2g shows the conformational changes along this direction (from the conformation with all three RBDs closed to a conformation with all three RBDs open).

Although the original study (which used STA and discrete classification) revealed only a fully closed state and a state with a single RBD open^22^, simultaneous opening of more than one RBD has been discussed in the literature^41^. However, considering the symmetry (C3) of the conformation with all three RBDs fully closed and our results showing that each of the three RBDs can undergo opening motion independently of the other two RBDs, we explored the possibility that a large number of the conformational models obtained by MDTOMO could be identical up to a rotation by 120° or 240° around the C3 pseudo-symmetry axis, which could have caused a difficulty in well separating the principal components by the PCA. Therefore, we designed a method to reduce the variability in order to simplify the PCA. This method is based on aligning the conformational models obtained by MDTOMO with respect to 5 conformations along the “RBD-A open” direction (Figure 2d), which describe opening of the RBD-A domain. Three out of the five conformational models used for this alignment are shown in Figure 2f. This method, which simplifies conformational variability analysis of the S protein, is explained next.

#### Simplifying PCA of conformational variability based on pseudo-symmetries of conformations

As introduced in the previous subsection, we reduced the number of principal components describing the variability due to RBD motions by aligning the conformational states obtained by MDTOMO with the states of a selected trajectory of a single RBD opening extracted from the initial PCA space (Figure 2d), which we here refer to as reference trajectory. The reference trajectory was selected as a five-state trajectory that corresponds to the most dominant single-RBD motion in the space determined by the first three PCA components, which is the opening of RBD-A (Figure 2d,f). The alignment was done by rotating the MDTOMO-derived conformations around the C3 pseudo-symmetry axis (here the z-axis) by multiples of 120° and matching the rotated conformations with the states on the reference trajectory.

More precisely, the set of 20,830 MDTOMO-derived conformational models was extended three times, by applying *n* × 120° rotation around the z-axis with *n* = {0,1,2}, resulting in 62,490 models (20,830 triplets, each triplet composed of the model rotated by *n* × 120°). Each of the three models in a triplet was compared by RMSD with the five models from the reference trajectory resulting in 15 RMSD values (five RMSD values per model of a triplet). The lowest value of the 15 RMSD values was identified and the associated rotation factor *n* was collected to determine the angle for the realignment of the MDTOMO-derived conformational model.

A new PCA was performed on the set of models obtained after the alignment, resulting in a new conformational variability space, described mostly by two principal components corresponding to a motion of RBD-A opening and a motion of NTD-B lifting, as shown in Figure 2e. Based on Figure 2e, it can be noted that the number of principal components necessary to describe RBD motions was reduced from 3 to 1. Also, in the new PCA space, the NTD motions are described by lower principal components (Figure 2c) than in the initial PCA space (components 2-4 in the new space vs. components 4-6 in the initial space).

The set of MDTOMO-derived conformational models aligned with the selected trajectory of single RBD opening (the trajectory shown in Figure 2f) was further analyzed using UMAP, which allowed a better separation of the different conformational states and identification of more populated distinct conformations.

#### UMAP representation of the conformational landscape simplified using pseudo-symmetries of conformations

The conformational landscape obtained after reducing the number of principal components to describe opening of RBDs (Figure 2e) shows a motion of RBD-A opening and a motion of NTD-B lifting. The set of aligned MDTOMO-derived conformational models was analyzed with UMAP to better separate different conformational states. The first two UMAP components are presented in terms of free-energy in Figure 3a. The conformational models and the density maps shown in Figure 3 are the conformational-model and subtomogram averages calculated from the four most populated regions (regions of lowest energy), manually selected and depicted in Figure 3a.

**Figure 3:**
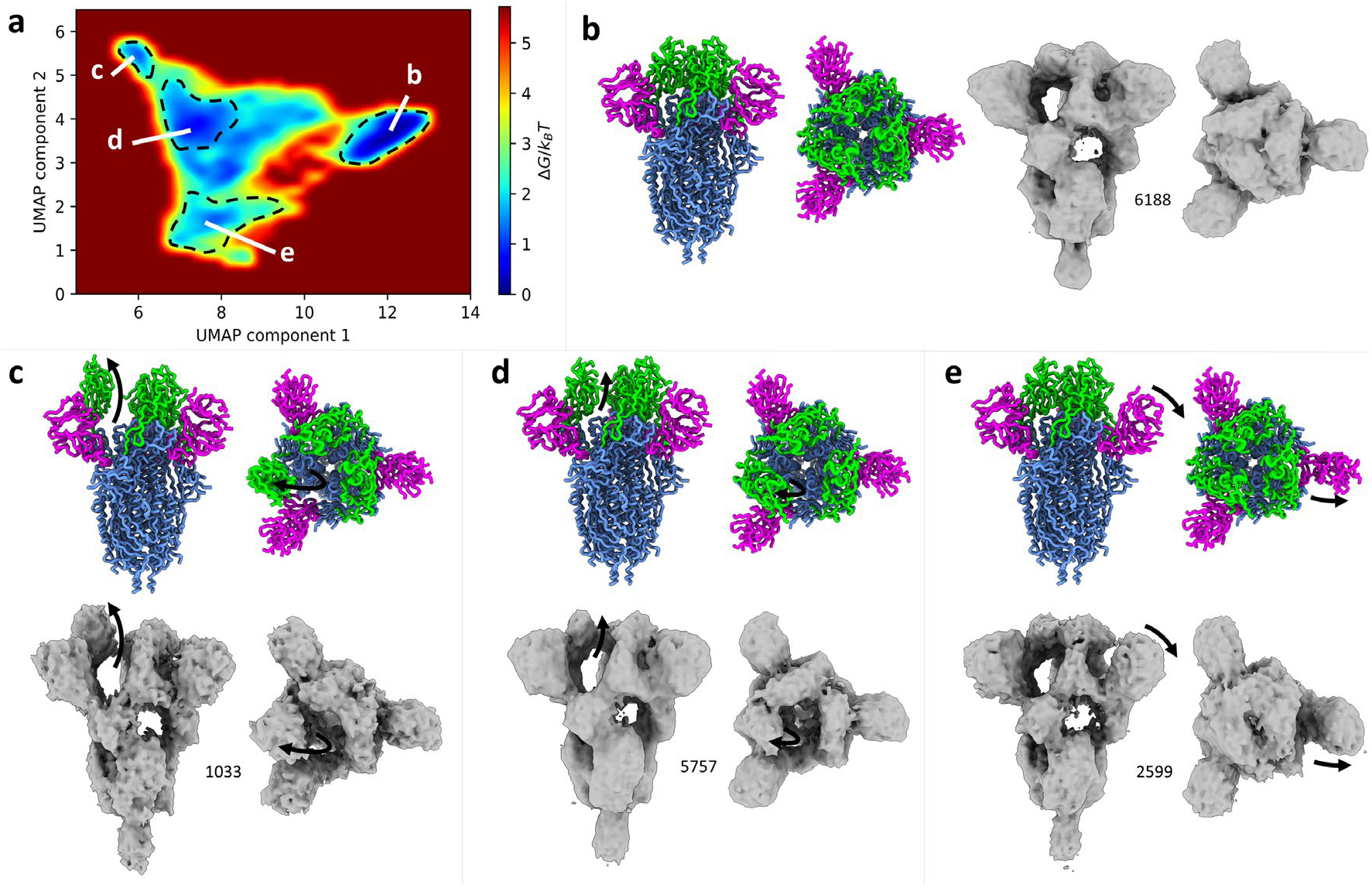
UMAP analysis of MDTOMO results with a cryo-ET dataset of SARS-CoV-2 S protein, after simplifying description of the variability due to opening of RBDs. (a) Free-energy landscape along the UMAP components 1 and 2. (b-e) Conformational-model and sub-tomogram averages from four manually selected regions in the free-energy landscape (denoted by b, c, d, and e). The colored model corresponds to the average of the conformational models in the selected region. The density maps are the averages of the subtomograms from the selected regions in the energy landscape. The number of the averaged particles is indicated near each map. Black arrows represent the conformational change from the initial structure used for the fitting within MDTOMO (PDB-6VXX, which is a conformation with all RBD closed).

The most populated region, designated as (b) in Figure 3a, corresponds to a conformer with all 3 RBDs closed. The corresponding conformational-model average and subtomogram average are shown in Figure 3b. The most distant low-density region from the region (b) along the first UMAP component, designated as (c) in Figure 3a, corresponds to a conformer with a single open RBD, i.e., RBD-A (Figure 3c). The central region, designated as (d) in Figure 3a, corresponds to intermediate states of a single RBD opening (Figure 3d). The bottom region along the second UMAP component, designated as (e) in Figure 3a, corresponds to a conformer with the NTB-B lifted (Figure 3e).

## Discussion

In this article, we presented MDTOMO, a novel approach to explore continuous conformational landscapes of biomolecular complexes by analyzing subtomogram datasets. MDTOMO uses efficient, coarse-grained and normal-mode-empowered MD simulations to extract the conformational states from subtomograms, which are typically averaged out during classical STA, and allows reconstructing low-dimensional (usually 2D or 3D) energy profiles of the structures from the experimental subtomogram data. MDTOMO is the first method for analyzing subtomograms using MD-based flexible fitting.

MDTOMO starts by extracting an atomic model from each individual subtomogram, by NMMD flexible fitting of a given reference atomic model into the subtomogram. The extracted atomic models are then projected onto a low-dimensional space, which can be considered as an essential conformational space. In this space, closer points represent more similar conformations, and denser regions of points represent the regions containing more likely or more stable (low-energy) conformations in the given dataset. The density of points in this space can be interpreted in terms of the free energy of the complex or by computing and analyzing the averages of the subtomograms and their associated atomic models in the regions of the space characterized by the lowest free energy of the complex.

In this article, we showed that MDTOMO produces expected results using synthetic data (ABC exporter). Also, we showed that the results produced using experimental SARS-CoV-2 spike protein data are coherent with previously published results regarding the existence of conformations with all three RBDs closed and those with a single RBD open^22^. Additionally, they revealed the existence of gradual conformational transitions related to the opening of single RBDs, a lifting motion of a single NTB, and additional conformations with more than one fully or partially open RBD.

Standard methods use the correlation coefficient as the metric for alignment and classification of subtomograms and require a compensation for the missing wedge (e.g., constrained correlation coefficient^42^) because of a sensitivity of the correlation coefficient to anisotropic deformations of subtomograms caused by the missing wedge. MDTOMO was able to produce expected results with synthetic and experimental data without compensating for the missing wedge, which can be explained by the fact that the NMMD flexible alignment in MDTOMO is not only driven by the force that depends on the correlation coefficient (derivatives of the correlation coefficient with respect to the atomic positions), but also by the classical MD-based force field (Eq. 1 and Eq. 4 in Vuillemot et al (2023)^13^). The weight given to the potential defined by the correlation coefficient, with respect to the classical MD potential, is a free parameter (the so-called force constant; *k* in Eq. 1 in Vuillemot et al (2023)^13^) whose value is set so as to drive the fitting toward the conformation in the subtomogram by preserving structural constraints, which reduces the risk of overfitting the noise and the missing-wedge-induced deformations in the subtomograms.

Compared to the majority of other methods for analyzing conformational flexibility in single particle cryo-EM datasets (e.g., 3DVA^10^ in cryoSPARC^43^ or cryoDRGN^11^) or cryo-ET datasets (e.g., Tomoflow^18^), MDTOMO uses an available atomic structure, whose coordinates are displaced (by performing NMMD simulation) to match the conformations in the subtomograms. Such atomic structure is not available for all datasets, but those datasets for which an atomic structure is available can efficiently be analyzed using MDTOMO. Indeed, MDTOMO incorporates structural constraints into the analysis via MD simulation, which is particularly important for analyzing cryo-ET subtomograms that suffer from noise and missing-wedge-induced deformations, as these structural constraints limit the motions to those that are feasible (underlying biochemical processes of the complex) and, therefore, they reduce the risk of overfitting mentioned previously. Also, the use of an available atomic structure in MDTOMO allows obtaining a conformational landscape in which the conformational transitions can be visualized directly on the atomic structure, contrary to the majority of other methods that visualize the conformational transitions on density volumes.

Another method that can use an available atomic structure to analyze conformational flexibility in cryo-ET datasets is HEMNMA-3D^19^. More precisely, HEMNMA-3D can use either an available atomic structure or an available density volume (e.g., subtomogram average or cryo-EM map)^19^, which is flexibly fitted into subtomograms by solely using normal mode analysis. Normal modes are useful for their simplicity and computational efficiency. However, they tend to induce distortions of the structure in case of large amplitudes of conformational changes. Also, they restrict the conformational exploration to a set of collective motions that must be defined in advance and that may be insufficient to describe the full range of conformational changes. MDTOMO overcomes these limitations by employing coarse-grained MD simulations to allow a full exploration of the conformational landscape without inducing structural distortions, and by empowering these MD simulations using normal modes to make simulations fast.

The MDTOMO approach requires running one MD simulation per subtomogram, which can be computationally expensive. The MDTOMO software is parallelized so that multiple simulations can run on multiple CPU cores simultaneously, with one simulation running only on one CPU core. Furthermore, the use of coarse-grained simulation coupled with normal-mode-based acceleration allows performing MDTOMO in a reasonable amount of time. The total execution time on 8 nodes of 2×20 Intel 6248 2.50 GHz for the two presented datasets is shown in Table 1. However, it is important to note that the computational cost scales with the molecular weight of the complex (the higher the number of atoms, the higher the cost of the MD simulation).

**Table 1:**
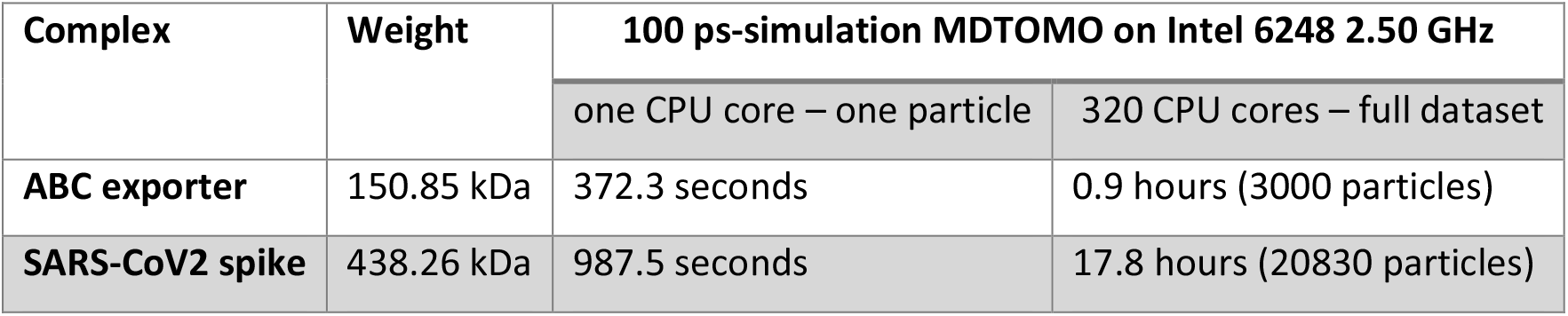
Execution time of MDTOMO for the two datasets analyzed in this article.

## Data Availability

The original contributions of the study presented here are included in the article files. The software code of MDTOMO will be publicly available on Github (https://github.com/scipion-em/scipion-em-continuousflex) and will also be part of the open-source ContinuousFlex plugin of Scipion. All questions regarding the software or data availability can be addressed to the corresponding author.

## Acknowledgments

We acknowledge the support of the French National Research Agency - ANR (ANR-19-CE11-0008 to SJ and IR and ANR-20-CE11-0020-03 to SJ); the cooperation between the CNRS and the University of Melbourne (The Melbourne-CNRS Network, PRC 2889 to SJ and IR, CNRS 80 Prime to SJ, joint PhD scholarship to RV); The University of Melbourne start-up fund (to IR); and access to HPC resources of CINES and IDRIS granted by GENCI (A0100710998R, A0100710998, A0070710998, AP010712190, AD011012188 to SJ). We thank Beata Turoňová (Max Planck Institute of Biophysics, Frankfurt am Main, Germany) for providing the subtomograms and the initial rigid-body alignment parameters that were used in the SARS-CoV-2 S protein experiment described in this article (from the dataset available in the EMPIAR database under the code EMPIAR-10453) as well as for critical reading of our manuscript and discussing the results that are presented here.

## Author Contributions statement

**Rémi Vuillemot**: Conceptualization, Methodology, Software, Investigation, Validation, Writing-Original draft preparation. **Isabelle Rouiller**: Validation, Writing - Review & Editing, Funding acquisition. **Slavica Jonic**: Conceptualization, Methodology, Investigation, Validation, Writing - Review & Editing, Project administration, Funding acquisition.

